# Phenology underlies apparent urbanisation effects on avian malaria in juvenile songbirds

**DOI:** 10.1101/2025.06.23.661047

**Authors:** Davide M Dominoni, Bedur Faleh Ali Albalawi, Magdalena Mladenova, Barbara Helm, Francesco Baldini

**Author notes:** these authors contributed equally (joint first authors) (Davide M Dominoni, Bedur Faleh Ali Albalawi).

## Abstract

Urbanisation can modify species interactions, including those between parasites and their hosts. In birds, urbanisation can either increase or decrease avian malaria infection, depending on host species, parasite or study location. However, temporal coordination between parasites and hosts, which may impact infection outcomes, has not been studied in urban ecology. To fill this gap, we collected blood samples from wild blue tit nestlings (Cyanistes caeruleus) in urban and forest habitats to examine how their hatch dates affected the prevalence and intensity of malaria infection. To separate parasites and quantify parasite load, we newly developed a species-specific qPCR assay. We found that Leucocytozoon prevalence was strongly affected by urbanisation, but effects depended on study year. This was driven by hatch date: nestlings that hatched earlier in the spring had a lower probability of being infected, independent of habitat type. In the few heavily infected nestlings, intensity of infection was associated with low body weight, suggesting fitness effects of infection. These results highlight the importance of breeding early to avoid early-life infection with malaria parasites, and that apparent urbanisation effects on infection arose from phenological differences between urban and forest habitats. Underappreciated phenological changes may underline other ecological effects of urbanisation.

## Introduction

As human populations continue to grow and cities expand into previously undeveloped areas, wildlife encounters with humans and human activities become more frequent [1,2]. This often leads to a restructuring of species composition and biodiversity via changes in several ecological processes, including species interactions [3]. An example of species interactions that may be affected by urbanisation are those between wildlife and their pathogens [4–7]. Evidence suggests that urban areas act as hotspots for the emergence of zoonotic diseases [5,8,9], including vector-borne diseases such as West Nile virus, Dengue fever, and Lyme disease [10–12]. These diseases can also be transmitted from wildlife to humans, thus posing risks to public health [13,14]. The anticipated surge in urbanisation worldwide is posed to further exacerbate disease risks [15,16], requiring a better understanding of host-parasite interactions.

Avian malaria parasites have been of central interest in the ecological literature of host-parasite interactions and vector-borne diseases over recent decades [17]. Identified in bird species around the world, it can pose significant threats to avian biodiversity, particularly for species in isolated island ecosystems that have not previously been exposed to the disease [18]. Its wide distribution also enables comparative studies across geographical areas and ecological contexts, including urban environments [17,19]. Avian malaria is caused by parasites of the genera *Plasmodium, Haemoproteus* and *Leucocytozoon*, which are transmitted through the bite of insect vectors [20]. These vectors are usually Culicidae mosquitoes in the case of *Plasmodium* species, *Culicoides* biting midges for *Haemoproteus* species and *Leucocytozoon caulleryi*, and *Simulium* blackflies for other *Leucocytozoon* species [21]. These parasites infect host blood cells (mainly red cells, but in case of *Leucocytozoon* also the white blood cells), tissues and other major organs such as liver, brain, heart and lungs [20].

The effects of haemosporidian infections on their host are complex as symptoms in birds can range from undetectable or minor illness to severe disease and death. In many cases, infected birds may show no visible symptoms but can still spread the disease through their vectors [20], yet infections can also affect various host traits. Examples include parental effort [22,23], body condition and body mass [24,25], global gene expression [26] as well as specific physiological processes such as iron balance [27] and telomere loss [28,29]. In turn, these changes have been suggested to negatively affect fitness-related traits, such as reproductive success [29–31] and survival [28,32,33], potentially leading to population decline [34]. Conversely, several studies have pointed to opposite or absent links between malaria infection and fitness [35]. Effects of malaria often depend on the parasite strain [36], on the species, population, age, sex or nutritional condition of the host [29,37,38] and on environmental conditions (e.g., food availability) during both growth and adulthood [39,40]. Several studies have also suggested that evolution of resistance in response to malaria infection should be more common than currently appreciated [18,41].

Studies of the impact of urbanisation on the prevalence of haemosporidian parasites showed mixed results. The relationship between urbanisation and malaria infection is context-specific and depends on several factors ([42–47], reviewed in [48]), including effects of year of study, latitude, city, and habitat type of surrounding rural areas (e.g., desert vs tropical rainforest). An overlooked factor that may modulate the impact of urbanisation on malaria parasite prevalence is phenology. Phenology, which affects disease patterns via fluctuations in abundance and condition of hosts, vectors and parasites [49–51], can differ between habitats [52–54]. The emergence and transmission of avian malaria in wild populations is strongly seasonal [55–58], partly due to seasonal changes in composition and abundance of vectors [56,59,60] and to seasonal relapse of parasites in birds [61]; additionally, year-to year variation in malaria prevalence is common [62,63]. Mosquitoes and other malaria vectors, such as midges and black flies, breed in stagnant or flowing water, and their populations can fluctuate depending on seasonal and yearly environmental conditions, such as temperature and rainfall [64,65]. During rainy seasons, when river flows and stagnant water reserves expand, vector populations can increase, thus potentially escalating the transmission of avian malaria. Conversely, in dry seasons or cold winters when vector populations decrease, the transmission of avian malaria is usually low [64]. In turn, aspects of phenology may differ between urban and non-urban habitats. One example are urban heat islands - areas in the city where temperatures are higher than in surrounding rural areas [66]. Higher urban temperature may increase the winter survival of vectors and advance both vector and bird phenology [67,68], further extending the transmission season of the disease [69]. The relative seasonal timing of host life-history stages and vector emergence may be affected by urbanisation, too, for example because species differ in response to ambient temperature [70,71]. Thus, urbanisation can potentially alter the seasonal patterns of avian malaria transmission and modulate its impact on urban bird populations.

Importantly, effects of many diseases also differ depending on host condition, physiology, and the timing of host, vector and parasite life-cycle stages in relation to each other [50,51]. For example, reproductive stages can be particularly sensitive to pathogens due to seasonal relapse in adults and developmental processes in offspring [72,73]. Phenological contributions to the prevalence of patent avian malaria infection (when parasites can be detected circulating in the blood rather than dormant in tissues) can be particularly well studied in nestlings, whose time of exposure had only started with hatching. Being naïve to infections, the nestlings’ host response is driven by innate immune responses directly linked to the physiological effects of the parasite (e.g. anaemia, tissue damage), rather than being shaped by acquired immune responses as in adults. Moreover, parasite intensity in nestlings can serve as a useful proxy for estimating the time since infection, as it reflects the progression of the acute phase [74]. Immediate, intensity-dependent effects — such as reductions in body mass— can therefore provide a clear window into the health costs of infection. Additionally, nestling body mass is a commonly used predictor for a nestling’s future fitness [75].

Thus, the aim of the present study is to disentangle effects of phenology and urbanisation on malaria in nestling songbirds. We study prevalence and intensity of avian malaria in early postnatal life of blue tits (*Cyanistes caeruleus*), a species that has served as a major study system in urban and disease ecology [76–79]. The blue tit is found across Europe in a variety of habitats, including urban, suburban, and rural environments, where they showed distinct responses to urbanisation [80,81]. Seasonal patterns of avian malaria have been described in both the blue tit and its arthropod vectors [49,60,81]. In Scotland and northern England, malaria vector emergence usually occurs in mid-late May, with a peak abundance in July [60,82–84]. Conversely, in the same areas blue tit nestlings hatch on average in early-mid-May. In Scotland, blue tit nestlings have been found to be infected with both *Haemoproteus* and *Leucocytozoon* parasites, but not *Plasmodium* [60,81]. The blue tits’ well-described breeding biology and physiology, including in our study populations, can aid in the interpretation of data on avian malaria and its impact. Thus, we here examine how the prevalence and intensity of infection with malaria parasites varied between forest and urban blue tit nestlings in Scotland, while explicitly considering the birds’ breeding phenology.

To disentangle factors that might blur associations, we also introduce methodological improvements. Although detection of haemosporidian parasites in host blood is relatively easy using molecular techniques and/or microscopy [85,86] it imposes challenges. The nested PCR protocol (nPCR) developed by Bensch et al. (2000) [87], and modified later [86], has been widely used to detect the three genera of Haemosporidia and to assess the reliability and sensitivity of newly developed protocols [68]. Nevertheless, the nPCR and other PCR-based protocols often overestimate mixed infection because of cross-reactivity of genus-specific primers [88]. Indeed, in a previous comparison of urban – non-urban blue tits, in nine out of 12 samples that were positive for both *Leucocytozoon* and *Haemoproteus/Plasmodium* by nPCR, sequencing revealed that only one of the two parasites – either *Leucocytozoon* or *Haemoproteus* – was present [81]. Thus, the field is in demand of a new sensitive and accurate molecular technique to quantify intensity of, and more importantly distinguish between, parasite genera in avian blood. Here we will apply a novel quantitative PCR approach that can distinguish and quantify *Leucocytozoon* and *Haemoproteus* with high sensitivity and specificity, which is particularly important when assessing parasite prevalence and intensity in nestlings when parasite development in peripheral blood is at its early stages.

With this approach, we here compare malaria infection in forest and urban blue tit nestlings across multiple years during which the birds’ breeding phenology varied distinctly. Specifically, we address the following questions:

## 1. Is the prevalence of malaria parasites affected by urbanisation and spring phenology?

We hypothesise that urban and forest blue tit nestlings differ in the prevalence of malaria parasites. We further predict effects of phenology of host reproduction on prevalence. In arthropods, spring emergence typically responds stronger to temperature than vertebrate reproduction, but patterns differ between terrestrial and aquatic species, and can be more variable between years in birds than in insects [82,89,90]. We hypothesise that in Scotland, where blue tits can hatch before vector emergence, low overlap between nestling growth and vector presence will reduce the risk of malaria infection. Thus, early-hatched nestlings should show lower prevalence than late hatched nestlings. Whether these effects are similar in urban and rural blue tits depends on how urbanisation affects host and vector phenology. Thus, if blue tits from both habitats breed on average early, we expect little malaria prevalence in both populations. However, if urban and rural birds differ in reproductive timing, inter-annual variation in host phenology could modulate differences in their malaria prevalence, assuming that vector phenology does not vary in synchrony.

### 2. Does malaria infection intensity impact nestling body mass and fledging success?

We hypothesise that malaria infection will impact on nestling condition and survival prospects. Specifically, we predict that malaria infection will reduce body mass before fledging and the probability of a chick’s fledging, independently of hatching habitat, but depending on time since infection as estimated from malaria intensity.

## Material and Methods

### Field work

The study took place in three forest sites and two urban sites in Scotland where ∼500 nest boxes were monitored for research. The forest sites were: (i) the Scottish Centre for Ecology and the Natural Environment (SCENE) (56° 7’ N, 4° 36’ W), (ii) the Sallochy campsite (56° 7’ N, 4° 36’ W) and (iii) Cashel farm (56° 6’ N, 4° 34’ W). The urban sites were in Glasgow: (i) Kelvingrove Park (55° 52’ N, 4° 17’ W) and (ii) the Garscube Sports Complex (55° 54’ N, 4° 18’ W). Descriptions of study sites and sample sizes for each site can be found in Table S1. We worked at these sites between April and June for four consecutive years, 2016-2019. At the beginning of April boxes were visited once a week to monitor nest building, laying date, clutch size and the start of incubation. After the tenth day of incubation (first day of incubation was defined as when the last egg was laid), nests were visited every other day to estimate hatch date. When nestlings were found in a nest, we estimated whether these had hatched the same day (day 0) or the day before (day 1) based on their appearance, and thus a hatch date was assigned to every nest. On day 13 after the first egg hatched (referred to as sampling date), we randomly selected three to four nestlings per nest, weighed them, and collected a blood sample (30-50 µl) by puncturing their brachial vein. Nest boxes were checked after fledging to search for any dead nestling. Each nestling was assigned a fledging success of 0 if it died before leaving the nest, or 1 if it successfully fledged.

### qPCR-based quantification of malaria infection

#### DNA extraction

DNA was extracted from blood samples using a commercial kit (DNeasy whole-blood extraction kit, Qiagen) and eluted with 80μL of water. Successful DNA extraction was confirmed using a Nanodrop ND-1000 Spectrophotometer (Nanodrop Technologies Inc., Wilmington, DE) and tested against the presence of glyceraldehyde-3-phosphate dehydrogenase (GAPDH) bird reference gene, following the protocol by [91], using primer pair: forward (5’-TGTGATTTCAATGGTGACAGC-3’) and reverse (5’AGCTTGACAAAATGGTCGTTC-3’) and the following thermal cycling profile: 95°C for 10 minutes, 40 cycles at 95°C for 1 minute, 1 minute at 60°C and 1 minute at 72°C, and then a final extension of 5 minutes at 72°C. The PCR products were then run out on a 1% agarose gel stained with SYBRTM Safe DNA Gel Stain (ThermoFisher Scientific, California, USA) and visualised under ultraviolet light.

#### Sequencing Leucocytozoon and Haemoproteus mitochondrial genes

To develop a qPCR that could quantify either *Leucocytozoon* or *Haemoproteus* parasites in bird DNA without cross-reactivity, we designed genus specific primers. First, we detected *Leucocytozoon* or *Haemoproteus* infected birds using the nested PCR protocol by [92], with some minor modifications (30 seconds instead of one minute for the denaturation, annealing and elongation steps). Then we amplified the conserved region of the mitochondrial genome between *cytB* and *cox1* previously used to detect avian haemosporidian infections using the primer set 292F/631R following [92]. Briefly, PCRs were performed in 20 μL reaction mixture using 1xGoTaq G2 hot start green master mix (Promega, USA), 0.32μM of each primer, and two μL of DNA. Cycling conditions were as follows: 5 min at 94°C, followed by 35 cycles of 30 seconds at 94°C, 30 seconds at 52°C and 30 seconds at 72°C, and a final elongation step at 72°C for 10 minutes. Amplification was evaluated on a 1% agarose gel, purified and sequenced on both strands (Eurofins Genomics).

#### Designing haemosporidia genus-specific qPCR primers

Obtained sequences (GenBank accession numbers TBC) were aligned using Clustal Omega [93] to reference intergenetic space between *cytB* and *cox1* genes from several haemosporidians, including *Leucocytozoon fringilinarum* (KY653765.1), *Leucocytozoon majoris* (FJ168563.1), *Leucocytozoon dubreuili* (KY653795.1), *Haemoproteus coatneyi* (KT698210.1), *Haemoproteus sacharovi* (KY653811.1), *Haemoproteus tartakovski* (KY653810.1), *Haemoproteus. sp*. (KY653805.1), *Haemoproteus belopolskyi* (KY653790.1); *Plasmodium relictum* (KY653773.1), *Plasmodium elongatum* (KY653801.1), *Plasmodium lutzi* (KY653816.1), *Plasmodium unalis* (KY653814.1) and *Plasmodium vaughani* (KY653792.1). Based on the alignment, we selected primer sequences that were different between genera, but that were conserved within each genus (Figure S1 and Table 1).

**Table 1.**
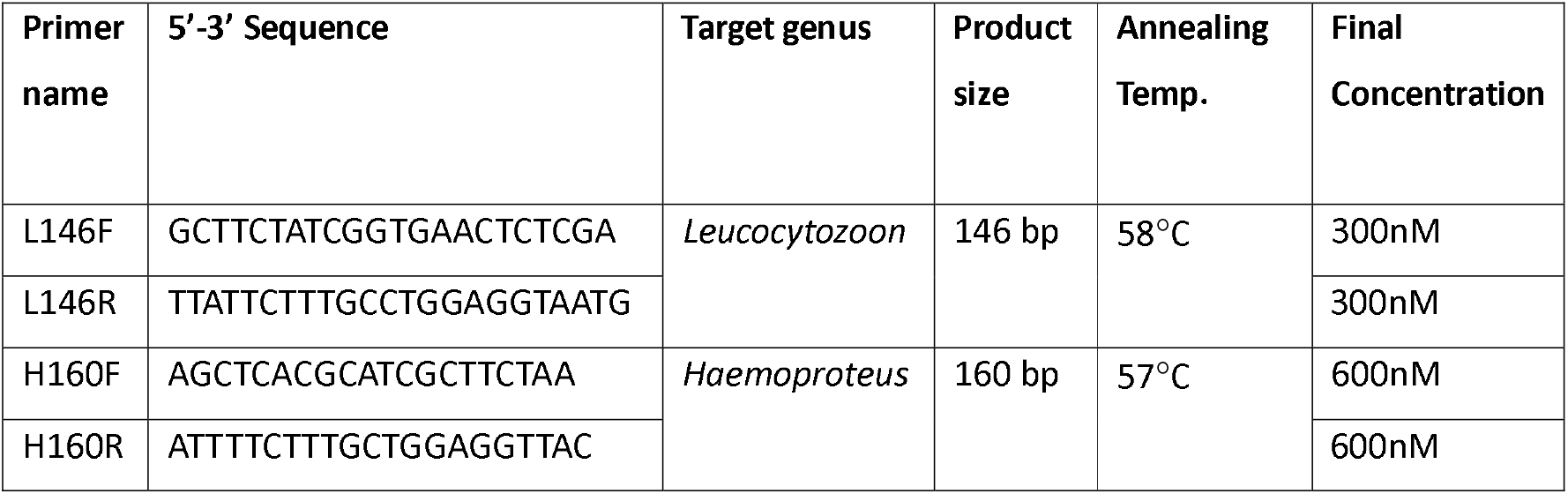
Haemosporidia genus-specific quantitative PCR primers.

#### Testing haemosporidia genus specific qPCR primers

To test the efficiency and specificity of the primers, amplicons were cloned (Invitrogen TA cloning kit) and qPCR tested on different numbers of plasmids from each target gene and two negative controls: uninfected bird sample (to ensure that primers are not amplifying bird genes) and distilled nuclease-free water. Samples were run in two technical replicates (accepted only if no more than 0.5 Cycle threshold (Ct) difference between the replicates). Best primer concentration for L146F/L146R was 300nM each with a primer efficiency of 94.1%, and for H160F/H160R it was 600nM each with a primer efficiency of 106.6%. When a primer set was tested against a different genus, only the most concentrated standard (10^7^ gene copies) showed cross-reactivity, but as this was observed ∼20 cycles after the expected Ct for the specific target, it suggested high specificity of each primer set (Figure S2).

#### qPCR quantification of parasite infections

*Leucocytozoon* and *Haemoproteus* infections were quantified by qPCR using our genus-specific primers against *Leucocytozoon* or *Haemoproteus* individually (Table 1), and normalised against the bird reference gene using GAPDH-Forward 5’-TGTGATTTCAATGGTGACAGC-3’ and GADPH-Reverse 5’-AGCTTGACAAAATGGTCGTTC-3’ at 100nM final concentration following [91]. The total volume of each reaction was 15 μl and included 5μL of DNA diluted to 12ng/μl, primers (see Table 1), 7.5 μl of SYBR Select Master Mix (Applied Biosystems), and nuclease-free water volumes. Reactions were run in MicroAmp Optical 96-well plates (Applied Biosystems) on Stratagene Mx3000 (ThermoFisher). Cycling conditions were as follows: 15 min at 95°C, followed by 40 cycles of 30 seconds at 95°C, 30 seconds at specific annealing temperature (57°C for *Leucocytozoon*, 58°C for *Haemoproteus*, 60°C for GADPH), followed by a melting curve. Quality checks were performed on the data and any duplicates from individual birds whose differences exceeded >0.5 cycle threshold (Ct) value were repeated. Additionally, samples that showed low quantities of the bird GADPH reference gene, i.e. amplified after 29 Ct, were excluded from the analysis as potential false negatives following observations that target parasite genes in low parasitemic samples tend to amplify 10 Ct after the reference. Finally, the relative infection intensity of malaria parasites was calculated for each sample as: x = 2^-(parasite Ct – GAPDH Ct)^.

## Statistical analyses

### General approach

All analyses were conducted in R, version 4.1.2 [94]. All models were linear or generalized linear mixed models (LMM or GLMM, respectively), which we ran using the R packages *lme4* [95] or *glmm*.*tmb* [96]. Nestbox identity was always included as random factor to account for variation explained by the same nestbox hosting different broods in different years, and to account for family effects (siblings sampled in the same nest). We checked that models adhered to assumptions of normality and homoscedasticity of residuals using the R package *performance* [97]. We used the same package to check for collinearity of predictors based on variance inflation factor (VIF). All models adhered to assumptions and collinearity was an issue only for year and hatch date (see below, VIF for all other predictors < 2). Model selection was conducted only when testing the significance of interactions, using likelihood ratio tests. If an interaction was not significant, it was removed from the model, but we left all main effects in the model as they were all crucial to test our hypotheses. All explanatory variables were scaled and mean-centered (across years) to improve the interpretability and comparability of model coefficients [98]. For example, when hatch date was used as an explanatory covariate, we calculated mean-centered hatch date as the deviation from the grand mean hatch date of all years, which was day 137 (May 17^th^), and then scaled it.

We first ran two models to describe variation in malaria parasite prevalence and hatch date, respectively, across years and habitats. These did not directly address our hypotheses but were instrumental to our understanding of the variation in malaria prevalence and timing of reproduction across years. In addition, hatch date and years caused collinearity problems, so they could not be included in the same models as covariates. We analysed the variation in malaria prevalence (0 = uninfected, 1 = infected) by fitting a generalized linear mixed model (GLMM) with a binomial error structure, using the function *glmer*. Habitat (a categorical factor with two levels: urban or forest), year, brood size and the interaction between habitat and year were included as fixed effects. We used the same approach, but applying a linear mixed model with the function *lmer*, to analyse variation in raw hatch date (un-centred). We then moved on to build models to answer our specific questions.

### Prevalence

we analysed the variation in malaria prevalence as function of hatch date and urbanisation with a binomial GLMM. Habitat, nestling hatch date (mean-centered across years) and brood size were modelled as fixed effects. We also included the interaction between habitat and hatch date to specifically test the hypothesis that the relationship between urbanisation and infection prevalence might depend on the phenological timing of the birds.

### Offspring fitness

To test for possible effects of malaria infection on nestlings, we first analysed the relationship between infection intensity and nestling weight. We used infection intensity rather than prevalence because higher intensity should be positively correlated to the number of days the animal has been infected for, and thereby the likelihood that it would cause effects on physiological state and condition [20]. For the nestling weight, we used a linear mixed model (LMM) with a Gaussian error structure, using the function *lmer*. Nestling weight was modelled as response variable, while infection intensity, nestling hatch date (mean-centered across years) and brood size were included as fixed effects. We also included the two-way interactions between intensity and habitat, intensity and hatch date, and hatch date and habitat. We repeated this same model using only birds that were heavily infected, which we defined as the upper 75% quantile (N = 11, Figure S3). We used a similar approach to analyse the relationship between malaria infection and a chick’s fledging success, but using a binomial GLMM with the probability of fledging (0 = dead, 1 = fledged) as response variable. However, because models that included infection intensity as predictor did not converge, we decided to use infection prevalence instead.

## Results

### Is the prevalence of malaria parasites affected by urbanisation and spring phenology?

*Haemoproteus* prevalence was very low in our dataset, as only 6 out of the 320 nestlings screened were infected with this parasite (1.8 %), equally found in urban (N = 3) and forest sites (N= 3). Conversely, *Leucocytozoon* prevalence was moderately high: 32.5 % of nestlings screened were infected with this parasite. In only two instances we detected co-infection. As *Haemoproteus* infection was very low and likely not playing a strong role in our system, we decided to focus only on *Leucocytozoon* for all the following analyses. *Leucocytozoon* prevalence differed between habitats, but depending on the year of study (Table S2): while prevalence was higher in forest than urban nestlings in 2016 and 2017, no difference between the two habitats was found in 2018 and 2019 (Figure 1A, Table S3). Similarly, mean hatch date over the four years spanned c. 20 days and differed between the urban and forest habitats (Fig 1b). However, the magnitude and direction of habitat differences depended on the year (Table S4). While forest nestlings hatched earlier than urban ones in 2016, 2017 and 2019, no difference was found in 2018 (Table S5).

**Figure 1.**
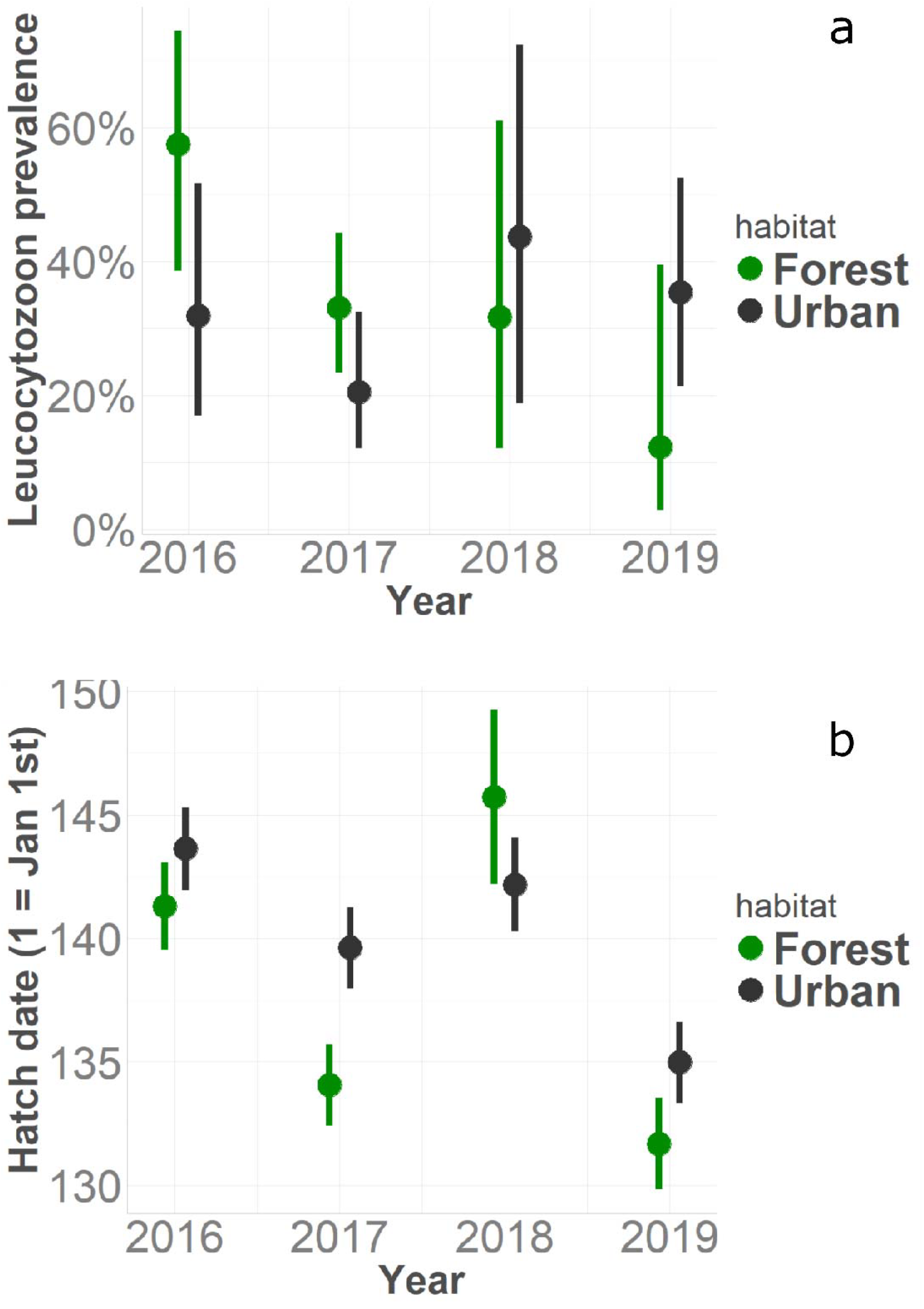
(A) Variation in *Leucocytozoon* prevalence in relation to habitat (forest vs urban) and year. (B) Variation in the date of nestlings hatching in relation to habitat (forest vs urban) and year of the study. Figures show predicted means ± confidence intervals.

The next step in our analyses was to test whether year-dependent differences in prevalence could be explained by nestling hatch date. As predicted, we found that *Leucocytozoon* prevalence did not differ significantly between urban and forest sites but was positively associated with hatch date (Table S6, Figure 2). All other variables included in our model did not affect *Leucocytozoon* prevalence (Table S6).

**Figure 2.**
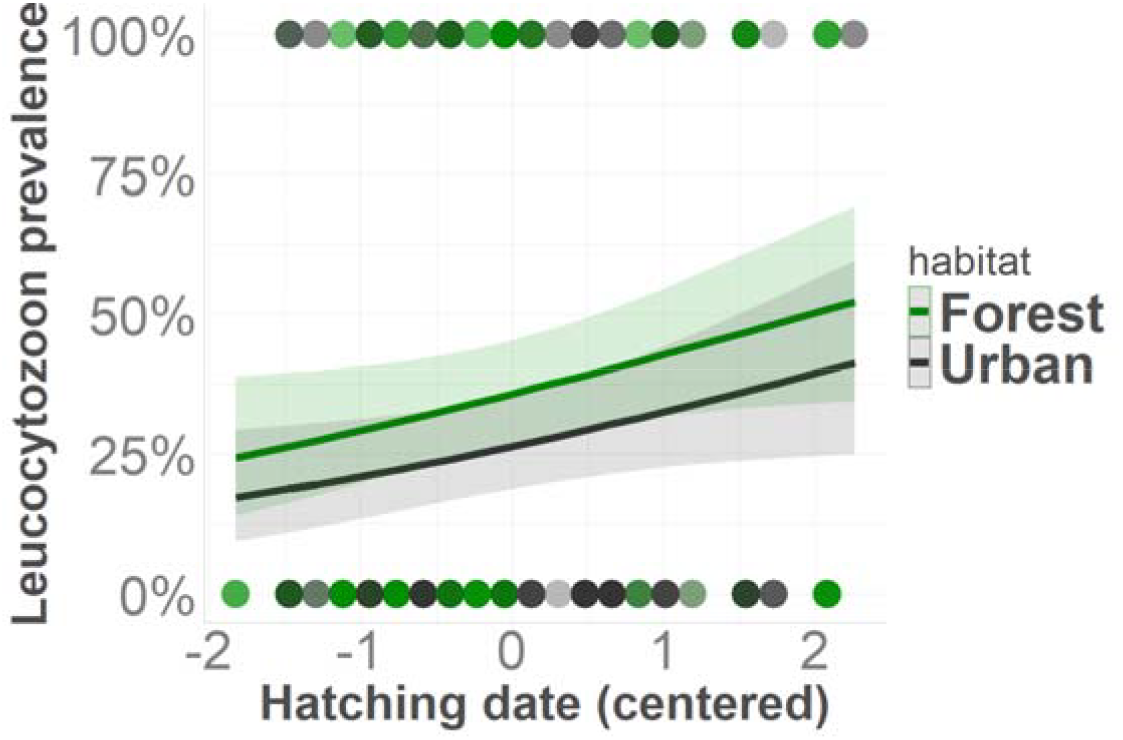
Variation in *Leucocytozoon* prevalence in nestling blue tits is positively related to the date of hatching. Nestlings hatched later in the spring have a higher probability to be infected with *Leucocytozoon*. This effect was independent of habitat. Line represents estimated means from Binomial GLMM, shaded area depicts confidence intervals. Points represent individual birds.

### Does malaria infection intensity impact nestling body mass and fledging success?

We found a strong and significant effect of hatch date on body weight, independent of habitat: both forest and urban nestlings were heavier later than earlier in the season (Figure 3a and Table S7). Moreover, forest nestlings were heavier than urban nestlings, but nestling body mass was unrelated to brood size (Table S7). Across all infected nestlings, body weight was not affected by *Leucocytozoon* infection intensity (Table S7). However, infection intensity was heavily skewed, with most birds showing intensity close to 0 and few birds with much higher intensity levels, which were all forest birds (Figure S3). As these low-intensity nestlings were likely only recently infected, we ran a subsequent model using only highly infected individuals to predict nestling weight. We defined highly infected individuals as those with relative level of infection above the 75% quantile of the data. Using this subset of 11 birds, our model showed that nestling weight was significantly negatively related to infection intensity: as infection intensity increases, nestling weight decreases (Figure 3b and Table S8).

**Figure 3.**
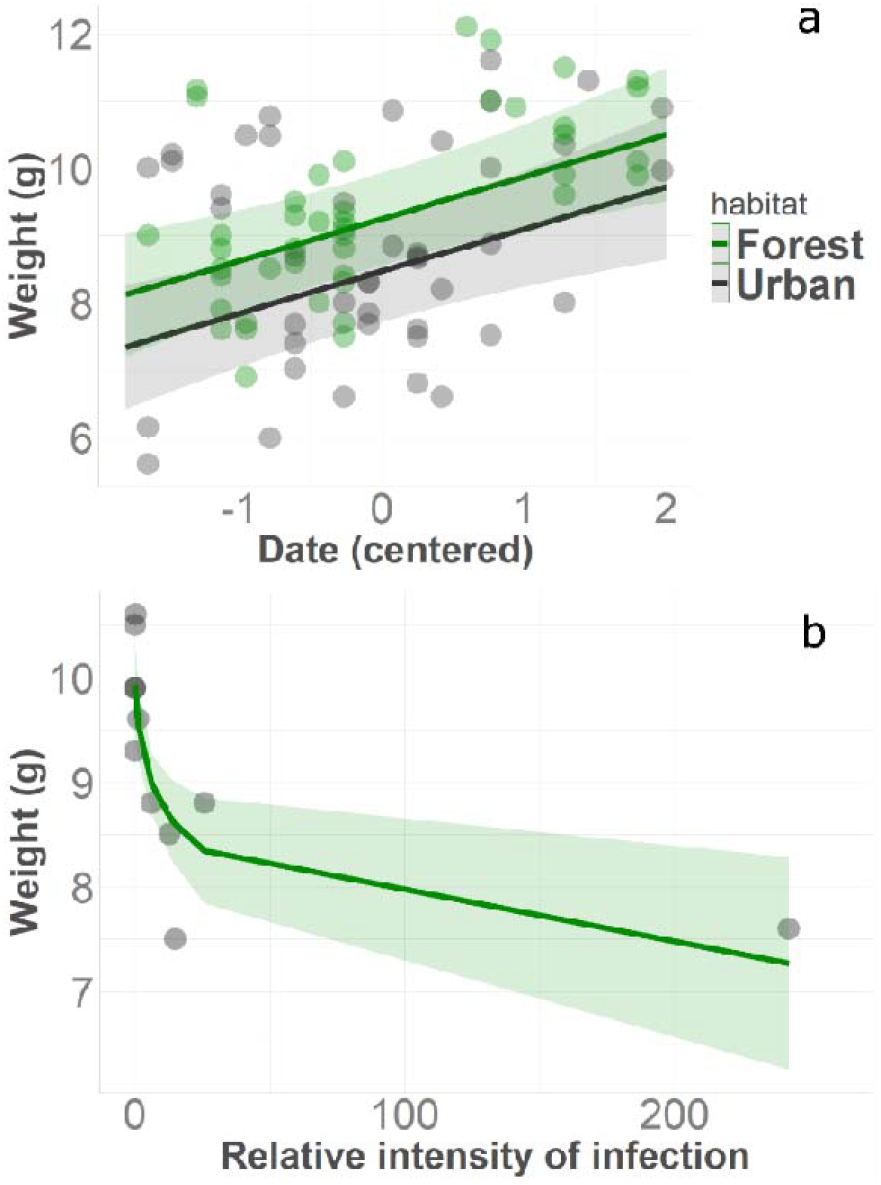
(a) Nestling weight is affected by season but not habitat. In both environments, nestling weight increases over the spring. (b) In the 11 highly infected individuals (which were all forest birds), nestling body weight is negatively related to *Leucocytozoon* infection intensity (parasite gene quantity relative to bird housekeeping gene). In both panels, lines are estimated means from Gaussian LMM, while shaded areas represent confidence intervals and dots represent raw data.

Across all nestlings, fledging success was significantly lower in the city than in the forest (Figure 4 and Table S9), and positively associated to nestling body mass (Table S9). *Leucocytozoon* prevalence, brood size and hatch date were not related to fledging success (Table S9).

**Figure 4.**
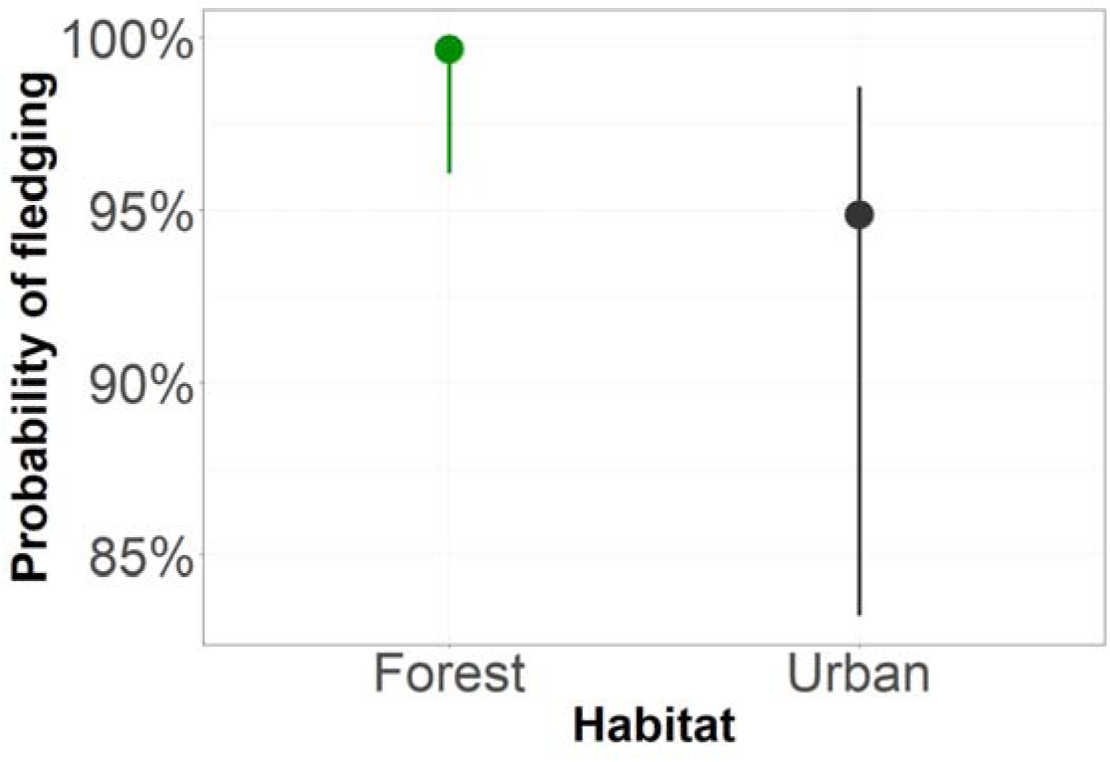
Urban nestlings have a lower probability of successfully fledging than forest nestlings. Points are estimated means over the four study years from a binomial GLMM, while error bars depict confidence intervals.

## Discussion

This is the first study, to our knowledge, that has explicitly investigated the interplay between phenology and urbanisation in determining avian malaria infection status. Our results provide evidence that the timing of hatching, rather than urbanisation, can explain variation in prevalence of a malaria parasite, *Leucocytozoo*n, in blue tit nestlings. In both urban and forest habitats, infection prevalence varied between years, but in no consistent manner. On average, nestlings that hatched earlier in the spring had a lower probability of being infected. This effect was quite strong: early in spring the probability of infection was as low as 20%, but later in the spring it rose to 60%. The phenology of hatching also influenced nestlings’ body weight, independently of habitat type. Nestlings born earlier in the spring were lighter than those that hatched later in the season, in contrast to previous studies in woodland passerines [99,100] and the widespread evidence that being early has fitness advantages [101]. Heavier nestlings were also much more likely to fledge. Low malaria infection intensity had no detectable effects on body mass of 13-day old nestlings. In the few nestlings that were found to be heavily infected, which were all forest birds, the intensity of infection was negatively related to body weight, suggesting that *Leucocytozoon* infection could potentially affect the health and fitness of blue tit nestlings. However, infection prevalence did not explain variation in fledging success across urban and forest nestlings.

Among the concise literature on avian malaria in the context of urbanisation, studies published to date have provided mixed results [48]. While generally avian malaria prevalence has been found to be lower in urban than non-urban birds of the same species, this pattern can vary depending on the species, life stage and local environmental conditions. For example, in house sparrows (*Passer domesticus*), studies conducted in Spain and USA found higher prevalence of *Leucocytozoon* in populations inhabiting rural habitats compared to urbanised areas [102,103], but no urban-rural differences were found in France [45]. In France, malaria infection in adult and nestling great tits (*Parus major*) was higher in urban than rural populations in Montpellier, but lower in urban than rural adults of the same species in Besancon and Dijon [46,104]. A study on five songbird species in an arid region of the southwest USA showed that adult birds generally exhibited lower *Haemoproteus* parasite prevalence in urban compared to rural populations, but species and sex-specific differences were found [47]. Our study confirms the mixed evidence. In some years malaria prevalence was higher in forest than urban nestlings, and in other years the pattern was reversed. Our analysis suggests that this pattern is likely explained by the fact that the time of hatching of nestlings in the two populations differed between the years, and that hatching time, rather than habitat type, drives differences in malaria prevalence between urban and forest birds. We thus assume that early hatching nestlings do not overlap much with the emergence of malaria vectors, thereby escaping infection.

Passerine bird phenology in temperate areas can vary considerably in time and space [105] and can be governed by both large-scale environmental variables, such as average spring temperatures [106,107], as well as by microenvironmental factors, such as the availability of insect prey [108]. Similarly, the phenology of the vectors that transmit malaria parasites can change across different years and over relatively small spatial scales. Thus, the mixed relationships observed between malaria prevalence and urbanisation may reflect variation in both reproductive phenology and vector emergence. Like bird reproduction, vector emergence is influenced by ambient temperature, but organisms vary in response to spring temperature. Thus, while terrestrial invertebrates generally advance phenology faster than vertebrates in warmer springs, these relationships are reversed in freshwater habitats, where the dipteran vectors develop [70,71]. Furthermore, different vector species might respond in distinct ways [65]. For instance, in our system, avian malaria is transmitted by blackflies, which breed in fast-flowing streams—habitats that may be less sensitive to temperature fluctuations and urbanisation than the small, temporary water bodies used by mosquito vectors. Such different developmental habitats can influence phenological patterns of vectors in an urbanisation context. Urban environments often provide artificial water reservoirs such as ornamental pond, fountains, gutters, and containers, which can serve as potential mosquito vector breeding grounds [69]. These microhabitats can sustain vector populations beyond the normal breeding season, potentially leading to more persistent transmission of avian malaria throughout the year in urban areas [69,109,110]. However, pollution and vector control activities in urban areas can considerably reduce larval numbers [111], and flowing waters – crucial for blackfly larval development - can be scarce in urban areas but more abundant in rural and agricultural areas. Future studies should investigate how urbanisation and seasonality may affect the abundance and phenology of different vectors.

In addition to hatching and vector phenology, seasonal changes of haemosporidian parasitaemia within blue tit hosts could also contribute to phenological effects on nestling malaria prevalence. After infection, birds enter the acute phase after which parasitaemia decreases drastically, known as the chronic phase (or latent stage). The chronic phase is no longer infectious, as parasites only persist in internal organs and not in peripheral blood [112]. However, when birds in temperate regions start breeding, an increase in parasitaemia is commonly observed [112]. This relapse is induced by changes in corticosterone and gonadal hormones, that are believed to directly trigger parasites in tissues [113,114]. Evidence of parasite relapse in *Haemoproteus* and *Trypanosoma* in blackcaps following increases in day lengths further suggest a suppressive role of melatonin, reducing relapses when nights are long [114]. Urban-related environmental factors known to affect phenology, such as artificial light at night, may thus also affect seasonal relapse of malaria parasites, e.g., via effects on melatonin or on the immune system [115]. In a system where most of the hosts have chronic infections, seasonal relapses that are timed with the vector’s peak in abundance are considered an adaptive strategy of the parasite to ensure transmission to the new-born susceptible birds [116]. Thus, synchrony between vector emergence and avian reproduction is key for parasite persistence across seasons [50], and these interactions could be facilitated, or reduced, by combinations of environmental change in urban habitats. We could not measure vector abundance and phenology at our sites, nor could we disentangle the effects of ALAN and other urban factors, which could have helped to explain our results. Future studies should integrate bird, vector and environmental data to gain a holistic understanding of how avian malaria infection varies in space and time with respect to urbanisation.

Our analyses also revealed a negative relationship between the intensity of *Leucocytozoon* infection and the nestlings’ body mass before fledging (i.e. when they were two weeks old), at least in those individuals that were heavily infected. High infection levels should reflect the time from first infection, and given that *Leucocytozoon* parasite life cycle is less than one week, it is likely that these nestlings were infected soon after hatching. However, malaria infection did not affect fledging success, which is more likely to be a function of poor diet leading to early nest mortality, especially in urban environments [117,118]. Nevertheless, since body mass before fledging is one of the most important predictor of future recruitment probability in small woodland passerines [75,119], our results point to potential long-term fitness consequences of early life infection with malaria parasites in these species, as suggested for great reed warblers (*Acrocephalus arundinaceus*) [28].

Molecular tools have significantly advanced avian malaria research by offering greater sensitivity and specificity than microscopy. Nested PCR (nPCR) has been instrumental in detecting *Plasmodium, Haemoproteus*, and *Leucocytozoon* [86,87], but it has limitations—namely, overestimating mixed infections due to primer cross-reactivity [88] and its inability to quantify parasite load. To address these issues, we developed a SYBR-based qPCR assay using species-specific primers for *Leucocytozoon* and *Haemoproteus*. This method requires two separate reactions, avoids the need for probes, and enables sensitive, cost-effective quantification. While validated only in Scottish samples, in silico analyses suggest broad applicability due to conserved primer regions across lineages. Compared to nPCR, our assay improves specificity by distinguishing single from mixed infections, detects down to 10^−5^ parasite gene copies per blood cell, and allows quantification. Other recent methods have tackled similar challenges: a genus-specific nPCR improves detection of mixed infections and phylogenetic resolution [120], and a multiplex PCR enables simultaneous detection of all three genera [121], though neither quantifies parasite load. Quantitative methods like a universal qPCR [122] and digital droplet PCR (ddPCR) [123] offer absolute quantification but cannot distinguish genera or detect mixed infections. A comparative study highlighted differences in detection rates across microscopy, multiplex, and nPCR, suggesting that combining methods may be optimal [124]. Future work should compare our qPCR with these approaches to evaluate its strengths and best-use scenarios.

Our study has some limitations. First, we sampled five populations in a small geographical area in and around Glasgow. Thus, it is unclear if our results can be generalized elsewhere. Second, as mentioned above, we do not have information on vector abundance and phenology in the different years and habitats, which does not allow us to verify our hypothesis that nestlings avoid overlapping with malaria vectors in years when they hatch earlier than average. This also prevents us from understanding how environmental conditions during spring, particularly temperature, can influence vector phenology, which might be particularly relevant in the context of climate warming, and how this might affect host-parasite interactions. Third, even though we sampled hundreds of birds over four years, *Leucocytozoon* prevalence was relatively low, especially in some years, which resulted in a small sample size for the fitness analyses. Moreover, *Haemoproteus* prevalence was extremely low and *Plasmodium* infection was never detected. The exact reasons still need to be clarified. For example, low or absent *Plasmodium* transmission might be a consequence of low abundance of *Culex* mosquito vectors; however this is less plausible for *Haemoproteus*, whose *Culicoides* biting midge vectors are highly abundant in the study sites. Alternatively, as the latent period of *Leucocytozoon* is much shorter (∼3 days) than in *Plasmodium* and *Haemoproteus* (∼10-14 days) [112], early sampling of nestlings at day 13 after hatching might have selectively favored the detection of the fastest parasite genus.

In summary, our study revealed previously unappreciated relationships between passerine reproductive timing, malaria infection and urbanisation. Our results highlight the importance of breeding early to avoid early-life infection with malaria parasites, at least in blue tits, and supports recent findings that infection can have a detectable impact on fitness-related traits. This has potential implications for climate warming, which is already leading many passerines to breed earlier. Whether insect vectors are also responding in the same direction, and to the same extent, is largely unknown, as long-term time series of vector phenology in relation to local environmental conditions are scarce. There is a clear research gap at the interface between bird and vector phenology, especially in an urbanisation context. Thus, we anticipate future studies to answer exciting questions on this topic.

## Acknowledgements

We thank Robyn Womack, Luke Woodford, Georgia Kirby, Heather Ferguson, and many master and undergraduate students who were involved with field sampling for this project. We thank the staff of SCENE, particularly Matt Newton and Hannele Honkanen, for hosting us and supporting our work. We thank Glasgow City Council and the Loch Lomond and the Trossachs National Park for giving us access to the urban and forest field sites, respectively. This research was funded by the University of Glasgow and by Funding through a Marie-Curie Career Integration Grant to BH [EC CIG (618578) Wildclocks]. BAA was funded by the Saudi Ministry of Education and by the University of Tabuk. FB was funded by AXA RF fellowship (14-AXA-PDOC-130), EMBO LT fellowship (43-2014), the Medical Research Council GCRF Infections Foundation Awards (MR/P025501/1) and the Academy Medical Sciences Springboard Award (ref:SBF007\100094).

## Notes

https://github.com/dmdominoni/urban_malaria/tree/main

### Competing Interest Statement

The authors have declared no competing interest.

