## supplementary tables and figures for "Phenology underlies apparent urbanisation effects on avian malaria in juvenile songbirds"

‡ co-corresponding authors

**Supplementary tables**

**Table S1.** Information on study sites. Habitat type, tree species composition, light and noise pollution are detailed for each site. Light and noise pollution were measured in May 2019. Light pollution was measured on cloudless nights using a photometer (LI-COR, LI-210 model). Noise pollution was measured twice in the morning between 9 and 11 am on consecutive days, randomizing the order of the nestboxes. Noise was measured using a sound pressure meter set on A frequency weighting (Precision Gold, N05CC model)

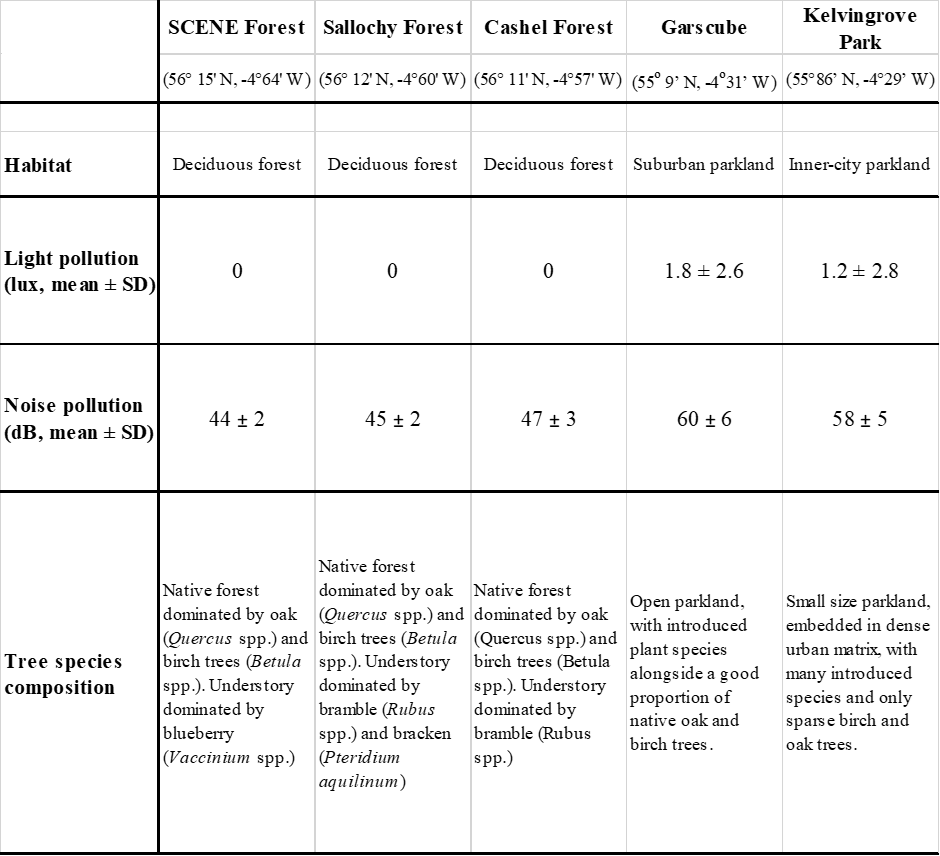

**Table S2.** The effect of year and habitat on *Leucocytozoon* prevalence. The model was a binomial GLMM with infection prevalence (0-1) as response variable, and habitat, year, brood size, nestling weight and the interaction year*habitat as fixed effects.

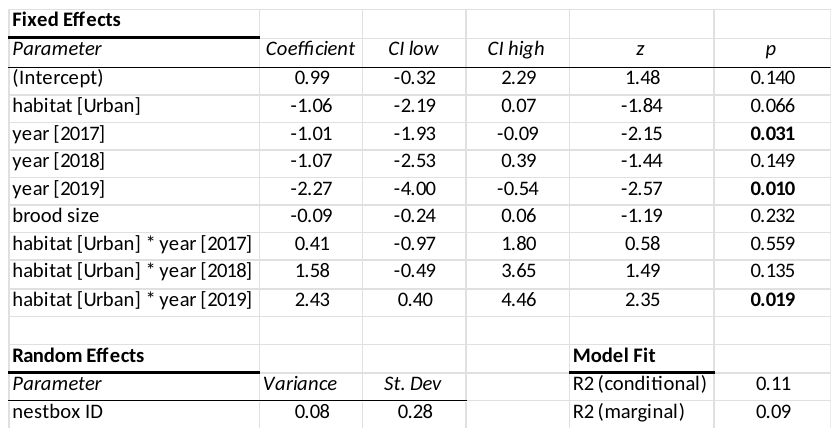

**Table S3.** Posthoc tests evaluating the significant interaction found in the model depicted in Table S2

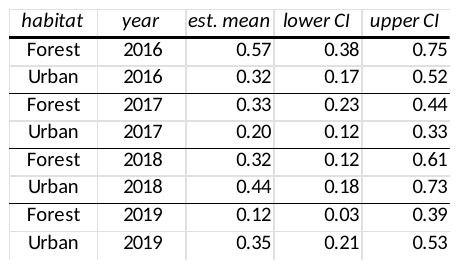

**Table S4.** The effect of year and habitat on nestling hatch date. The model was a Gaussian LMM with hatching date as response variable, and habitat, year, brood size, nestling weight and the interaction year*habitat as fixed effects.

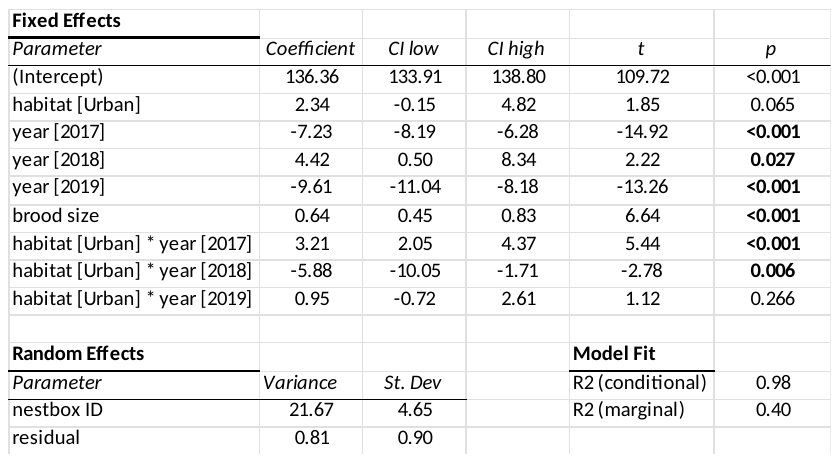

**Table S5.** Posthoc tests evaluating the significant interaction found in the model depicted in Table S4

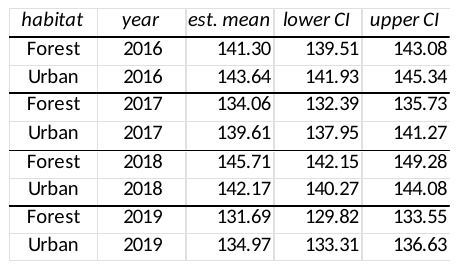

**Table S6.** The effect of nestling hatch date and habitat on *Leucocytozoon* prevalence. The model was a binomial GLMM with infection prevalence (0-1) as response variable, and habitat, hatch date, brood size, nestling weight and the interaction hatch date*habitat as fixed effects. This interaction was removed as it was not significant.

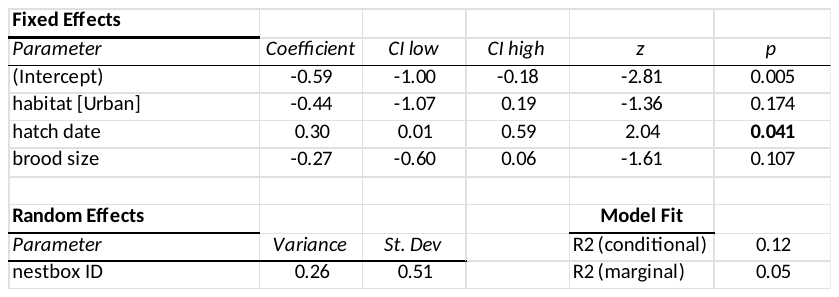

**Table S7.** The effect of nestling hatch date, habitat and *Leucocytozoon* infection intensity on nestling weight. The model was a Gaussian GLMM with nestling weight as response variable, and habitat, hatch date, brood size, *Leucocytozoon* intensity, as well as the interaction hatch date*habitat, as fixed effects.

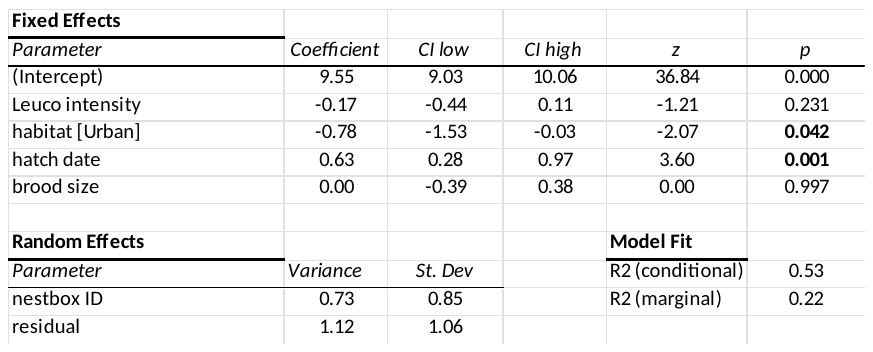

**Table S8.** The effect of nestling hatch date, habitat and *Leucocytozoon* infection on nestling weight, using a reduced dataset with only highly infected nestlings (which were all forest nestlings, so habitat was not included in the model). The model was a Gaussian GLMM with nestling weight as response variable and hatch date, brood size, and *Leucocytozoon* intensity as fixed effects.

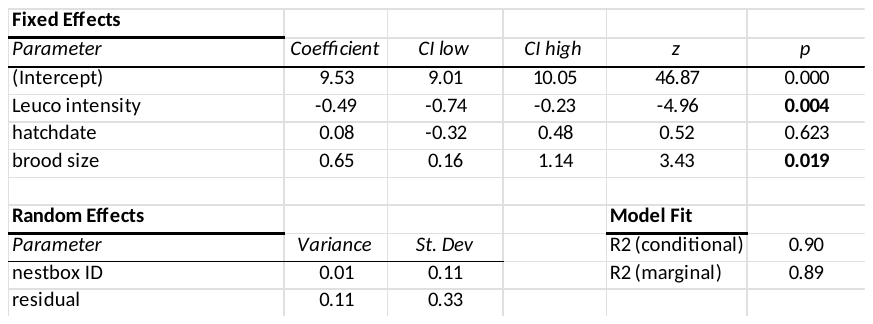

**Table S9.** The effect of nestling hatch date, habitat and *Leucocytozoon* infection on nestling fledging success. The model was a Binomial GLMM with nestling fledging success as response variable (0-1), and habitat, hatch date, brood size, centred date, *Leucocytozoon* prevalence, as well as the interaction hatch date*habitat as fixed effects. This interaction was removed as it was not significant.

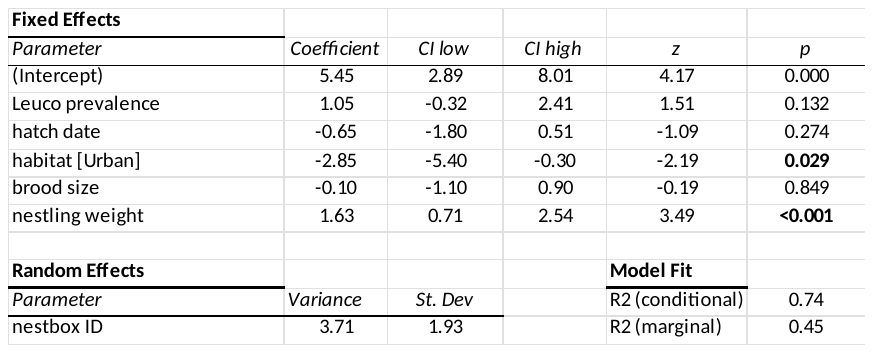

**Supplementary figures**

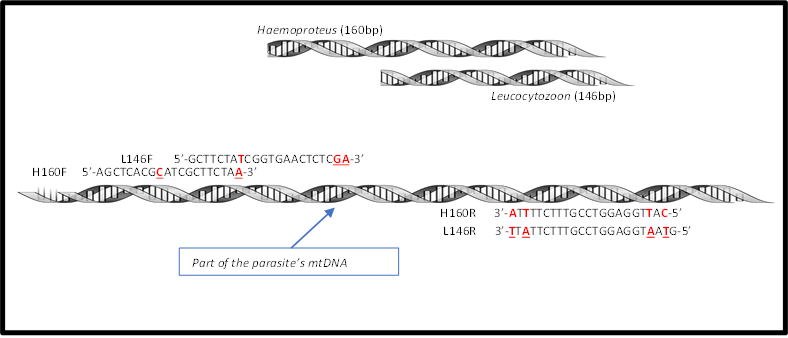

**Figure S1.** Schematic illustration of the designed *Haemosporidia* genus-specific qPCR primers.

L146F/L146R primer set amplifies a fragment (146bp) of the target gene of *Leucocytozoon* genera. H160F/H160R primer set amplifies a fragment (160bp) of the target gene of *Haemoproteus* genera. Underlined letters indicate the primer sequences that were not shared between the two genera.

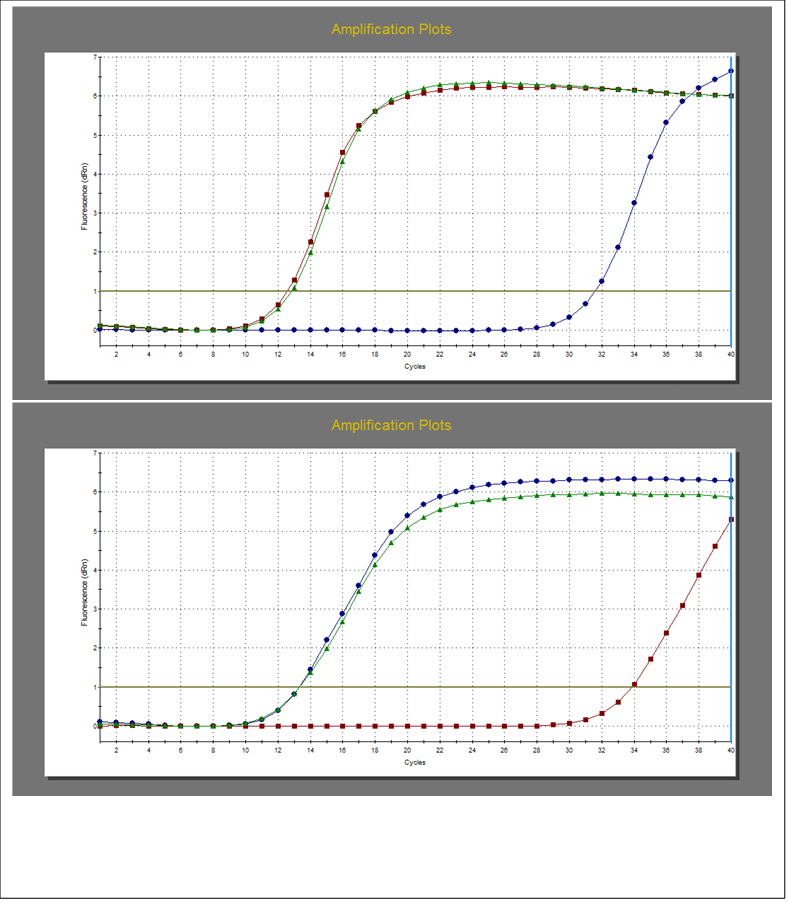

**Figure S2.** qPCR amplifications of three different plasmids (10^7^ gene copies) (green line with

triangles: mix of *(Haemoproteus* and *Leucocytozoon* plasmids), red line with squares: *Haemoproteus* plasmid, and blue line with dots: *Leucocytozoon* plasmid). A) shows amplifications by using *Haemoproteus* primers while B) shows amplification by using *Leucocytozoon* primers. The

horizontal line in the amplification plots indicated the fluorescence threshold.

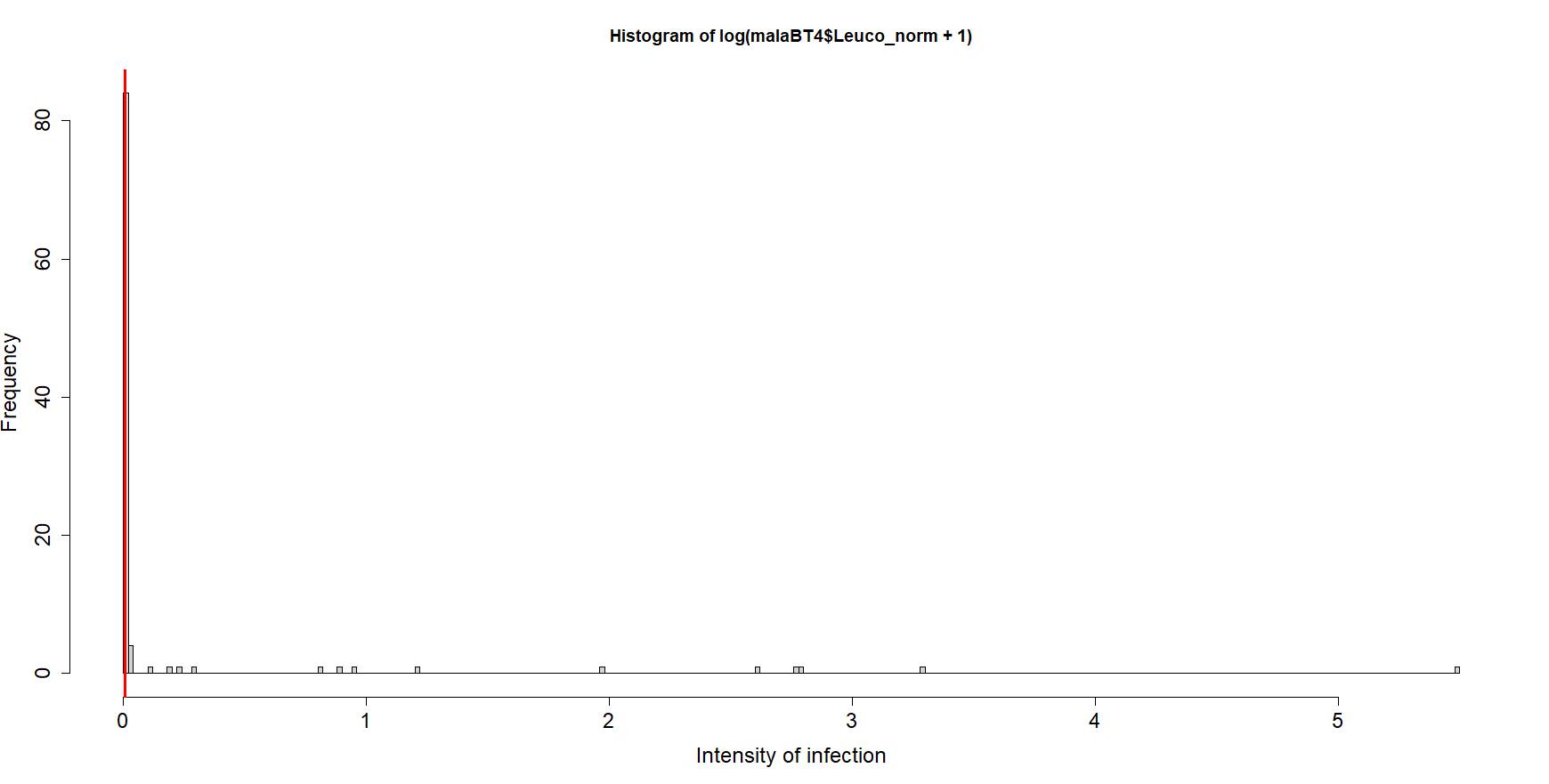

**Figure S3.** Histogram of relative infection intensity. Highly infected individuals were considered to be those above the 75% quantile of the data (0.005, depicted by vertical red line).
